# Linking UV absorption of DNA with base pair sequences

**DOI:** 10.64898/2026.05.05.722932

**Authors:** Luc Wieners, Martin E. Garcia

## Abstract

The absorption of ultraviolet (UV) radiation is the first step in a chain of events which can induce DNA damage associated with cancer and aging, yet the sequence dependence of UV absorption remains to be investigated. Here, we present a systematic study of UV absorption spectra of DNA based on all-electron Hartree–Fock calculations. We analyze all possible sequences up to four base pairs, as well as longer randomized sequences and genomic nullomers—motifs which are missing in a given genome. We observe a pronounced sequence dependence: cytosine- and guanine-rich motifs exhibit significantly enhanced absorption, whereas adenine–thymine-rich sequences absorb less in the mid-UV range. Most genomes are biased toward adenine–thymine-rich sequences which leads to decreased absorption of UV radiation by the sun. In addition, we introduce a computational framework enabling spectral calculations of large DNA and RNA fragments, opening the door to large-scale optical analyses.

---

The human genome contains around 3 billion base pairs which are the smallest units of information in DNA, each containing around 60 atoms. This information is continuously exposed to environments which can damage it, with estimates of around 10,000 instances of DNA damage occurring per cell each day [1]. Evolution has given rise to numerous DNA repair mechanisms that, in most cases, mitigate damage caused by environmental and endogenous agents [2]. However, in situations where DNA repair fails, the sustained DNA damage can lead to aging and cancer.

Among the environmental causes of DNA damage, ionizing radiation plays an important role, especially in the form of ultraviolet radiation due to everyday exposure to the sun. UV radiation can directly cause damage by forming pyrimidine dimers [3] which cause distortions to DNA structure.

To have UV radiation cause harm to organisms, several events have to take place. First, the UV radiation has to be absorbed by DNA. Then, this absorption needs to create a damage in DNA which happens with certain probabilities [4]. Finally, the corresponding damage has to survive DNA repair and cause a mutation that is harmful to the organism. In this work, we focus on UV absorption as the starting point of this chain of events.

Kufner et al. [4] investigated the sequence-dependent UV damage formation for DNA tetramers and hexamers. Additionally, there have been theoretical spectral investigations of nucleobases [5] and the AT dimer [6]. However, a quantitative understanding of how UV absorption specifically is related to DNA sequences in the genome is still to be investigated. In this Letter, we demonstrate using all-electron Hartree–Fock calculations that UV absorption in DNA is strongly sequence-dependent, with enhanced absorption in cytosine–guanine-rich motifs.

Ultraviolet (UV) radiation is conventionally classified into UVA (315–400 nm), UVB (280–315 nm), and UVC (100–280 nm) bands according to wavelength. Solar UVC is almost completely absorbed by the atmosphere and does not reach the Earth’s surface in significant amounts. Consequently, terrestrial UV radiation consists primarily of UVA and UVB, with UVA dominating due to its weaker atmospheric attenuation. However, DNA exhibits negligible absorption at wavelengths above 320 nm; therefore, we focus on UVB radiation in this study.

To obtain systematic insight into the UV absorption of DNA samples, we computed spectra for all possible sequences up to four base pairs, comprising 16, 64 and 256 structures for lengths with one to four base pairs, respectively. In addition, we analyzed longer sequences of eleven base pairs, including ten nullomers (DNA sequences which do not occur in the human genome) [7, 8] and ten of their synonyms sequences—distinct base-pair sequences encoding the same translation. As a reference, we also considered ten random DNA sequences.

DNA structures containing one to four base pairs comprise approximately 60, 125, 190, and 250 atoms, respectively, while sequences with eleven base pairs contain about 700 atoms. Fully quantum-mechanical calculations of absorption spectra become computationally prohibitive for systems exceeding 100–200 atoms. In such cases, QM/MM methods [9] that combine molecular mechanics (MM) [10–13] with a quantum-mechanical (QM) treatment [14, 15] of a selected (core) region are typically used. Here, however, our aim is to capture the absorption of the entire DNA structure rather than a localized region. We therefore adopt a fully quantum-mechanical treatment of the complete system, with reduced accuracy to maintain computational feasibility.

Our computational framework [16] is based on real-time time-dependent Hartree–Fock (RT-TDHF) and employs a minimal STO-3G basis, truncated long-range interactions, and the intrinsic approximations of the Hartree–Fock method. While this level of theory does not provide quantitatively accurate excitation energies, it yields a consistent description across the large set of structurally similar DNA sequences considered here, enabling robust identification of relative trends in UV absorption. The limitations arising from these approximations - including the neglect of electronic correlation and interaction truncation - are largely systematic, as the sequences share a common backbone structure, and can therefore be mitigated through comparative analysis over the full dataset. Additionally, electronic structure benchmarks using TDDFT (def2-TZVP, CAM-B3LYP) were performed for monomers, dimers, single nucleobases and nucleotides, the effects of water, ions and conformations were studied as well (see Supplementary material, Figures 2–7). Details on the parametrization and validation of our approach are provided in Ref. [16]. For discussions of systematic error correction in TDHF and TDDFT, we refer to references [17, 18].

For the computation of DNA absorption spectra, we construct sequences composed of the four nucleobases adenine (A), cytosine (C), guanine (G), and thymine (T). Atomic coordinates are generated using the nucleic acid builder in UCSF ChimeraX [19–21]. To account for the negative charges associated with the phosphate groups in the DNA backbone, additional electrons are included. Specifically, for a DNA sample with *n* base pairs– corresponding to *n −* 1 backbone linkages per strand–we add 2*n −* 2 electrons.

Following structure preparation, quantum-mechanical time propagation is performed using RT-TDHF (see Supplemental Material). The computational cost is dominated by matrix operations, particularly matrix exponentiation, and the repeated evaluation of the electrostatic potential during the time evolution of the electron density. We employ the Hartree–Fock method because it enables efficient electrostatic calculations for the spatially extended DNA structures considered here.

Within the Hartree–Fock formalism, the integrals required for the construction of the Fock matrix can be stored in memory (10^8^ *−* 10^9^ integrals for the systems considered here). Computational efficiency is further enhanced by truncating long-range interactions, which has little impact on RT-TDHF spectra [16].

Inclusion of all interactions remains feasible, albeit at a several-fold increase in computational cost depending on system size. Benchmark calculations for DNA dimers revealed only minor deviations from the truncated-interaction results, well below the uncertainties associated with the minimal-basis Hartree–Fock approximation. Overall, RT-TDHF enables a substantially faster construction of the Hamiltonian matrix, which constitutes the primary bottleneck in large-scale RT-TDDFT calculations [22].

Systematic errors can be reduced through empirical calibration procedures that are particularly effective for classes of structurally similar molecules [17, 18]. Such approaches typically employ a linear regression between experimental and theoretical peak positions. Here, we instead apply a wavelength-dependent linear correction over an interval, as our analysis focuses on the spectral shape in the UVB region rather than on individual absorption peaks.

Therefore, we use the experimental DNA absorption spectrum (absorbance vs. wavelength) between 220 and 320 nm as a reference [23], with particular emphasis on the UVB region near 300 nm. The corresponding theoretical reference is obtained from the average spectrum of all tetramers. Experimental absorbance spectra can be compared with theoretical absorption spectra since both describe the same interaction between light and matter; however, absolute intensities are not comparable.

Key features of the experimental DNA absorbance spectrum are a minimum near 230 nm, a maximum near 260 nm, and a decay to nearly zero signal at 320 nm. Additional reference values are provided by the ratios (*A*_260_*/A*_280_ *≈* 1.8) and (*A*_260_*/A*_230_ *≈* 2.0). To constrain the UVB region, we furthermore require the ratio between the spectral intensity at 300 nm and the maximum near 260 nm to be approximately 0.05.

Optimizing the scaling factor and offset yields the correction function *λ*_corr_ = 1.1 *· λ*_calc_ + 20 nm, which reproduces the experimental spectral characteristics discussed above. A sensitivity analysis of the wavelength correction equation is shown in Supplementary material, Figure 8. The corrected spectra of all 256 tetramers, their average, and the comparison with experiment [23] are shown in Fig. 1, including an enlarged view of the UVB region.

**FIG. 1.**
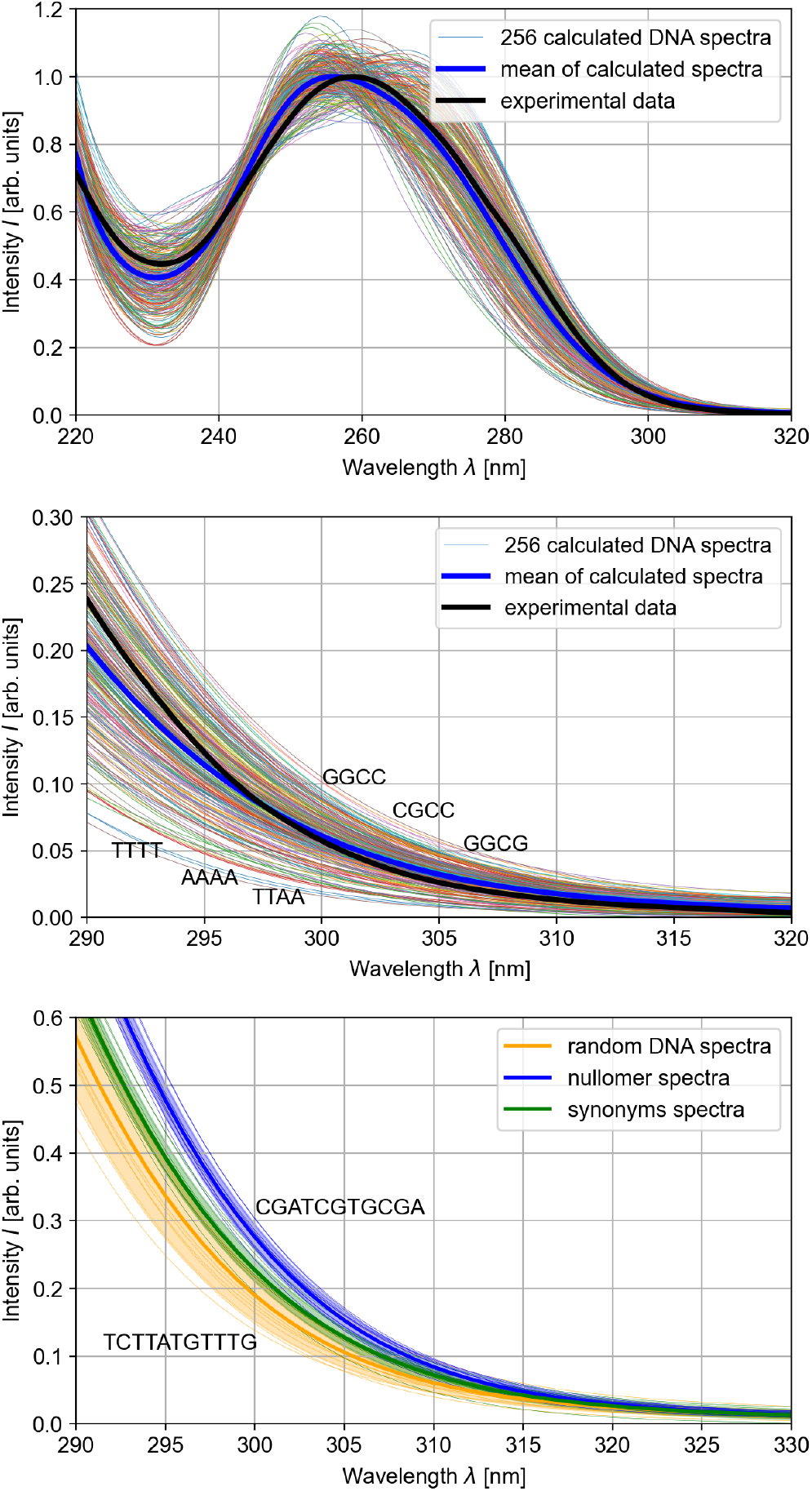
Top: The corrected spectra of the 256 possible DNA tetramers are shown as colored lines. Additionally, the mean of these spectra is depicted (blue) and compared to an experimental spectrum of DNA (black) [23]. Middle: The results from the upper panel are shown in the region between 290 and 320 nm with the sequences with the lowest and highest absorption highlighted. Bottom: The absorption spectra of the ten investigated nullomers (blue), their synonyms (green) and of ten random DNA samples (yellow) are shown. The individual spectra are shown as narrow lines, the averages as wide lines and the standard deviations as colored background.

We further investigated larger DNA structures by computing the absorption spectra of ten randomized eleven-base-pair sequences, as nullomers begin to emerge at this length in the human genome. The randomized sequences were generated such that their average base composition matches that of the human genome (*∼* 30% adenine, *∼* 30% thymine, *∼* 20% cytosine, and *∼* 20% guanine).

We considered the ten eleven-base-pair nullomers listed in Table 3 of Ref. [7], namely CGCTCGACGTA, GTCCGAGCGTA, CGACGAACGGT, CCGATACGTCG, TACGCGCGACA, CGCGACGCATA, TCGGTACGCTA, TCGCGACCGTA, CGATCGTGCGA, and CGCGTATCGGT. Notably, these sequences are enriched in cytosine and guanine, particularly in the form of CG motifs, despite the comparatively low GC content of the human genome (40.89%) [24].

Corresponding synonymous sequences were generated from the nullomers by replacing codons with synonymous alternatives that preserve the encoded amino-acid sequence. This is possible because the genetic code is degenerate, with 64 codons encoding only 20 amino acids. The nullomers were converted into synonymous DNA sequences by partitioning each sequence into four codons. Each codon was translated into its corresponding amino acid and subsequently replaced by the codon most frequently used for that amino acid in the human genome. Because a DNA sequence can be partitioned into codons in three different ways, depending on the reading frame, the resulting synonymous sequence depends on the chosen frame. In the present analysis, the reading frame begins at the first nucleotide. Since the final codon contains only two nucleotides, the third nucleotide was chosen according to the most probable codon consistent with the first two positions.

Analysis of the 30 longer DNA sequences (random, nullomer, and synonymous) reveals systematic differences in their UV absorption spectra (Fig. 1). The random sequences exhibit the largest variability in absorption near 300 nm, as reflected by the orange band indicating one standard deviation.

In contrast, the nullomers and their synonymous counterparts display considerably less variation, and the average nullomer absorption lies beyond the standard-deviation range of the random sequences. This enhanced absorption is likely attributable to the pronounced enrichment of cytosine and guanine in the nullomer sequences.

The average UV absorption of the synonymous sequences is significantly lower than that of the nullomers, but remains within the upper range of the random DNA samples. This suggests that the use of more common codons reduces susceptibility to UV radiation, yet does not fully compensate for the enhanced absorption of the nullomer sequences. Such incomplete compensation is expected, as the translation of nullomers into synonymous sequences offers only limited flexibility. In many cases, synonymous codons differ by only a single nucleotide, leaving substantial portions of the original sequence unchanged.

We next compare the UV absorption of all possible DNA monomers, dimers, trimers, and tetramers with their occurrence frequencies in the human genome (Fig. 2). For all sequence lengths, we observe a pronounced inverse correlation between genomic abundance and absorption intensity at 300 nm. The corresponding correlation coefficients are (-0.937), (-0.633), (-0.705), and (-0.692) for monomers, dimers, trimers, and tetramers, respectively. Consistently, the inverse absorption intensity exhibits positive correlations with genomic abundance, yielding coefficients of (0.857), (0.707), (0.745), and (0.721).

**FIG. 2.**
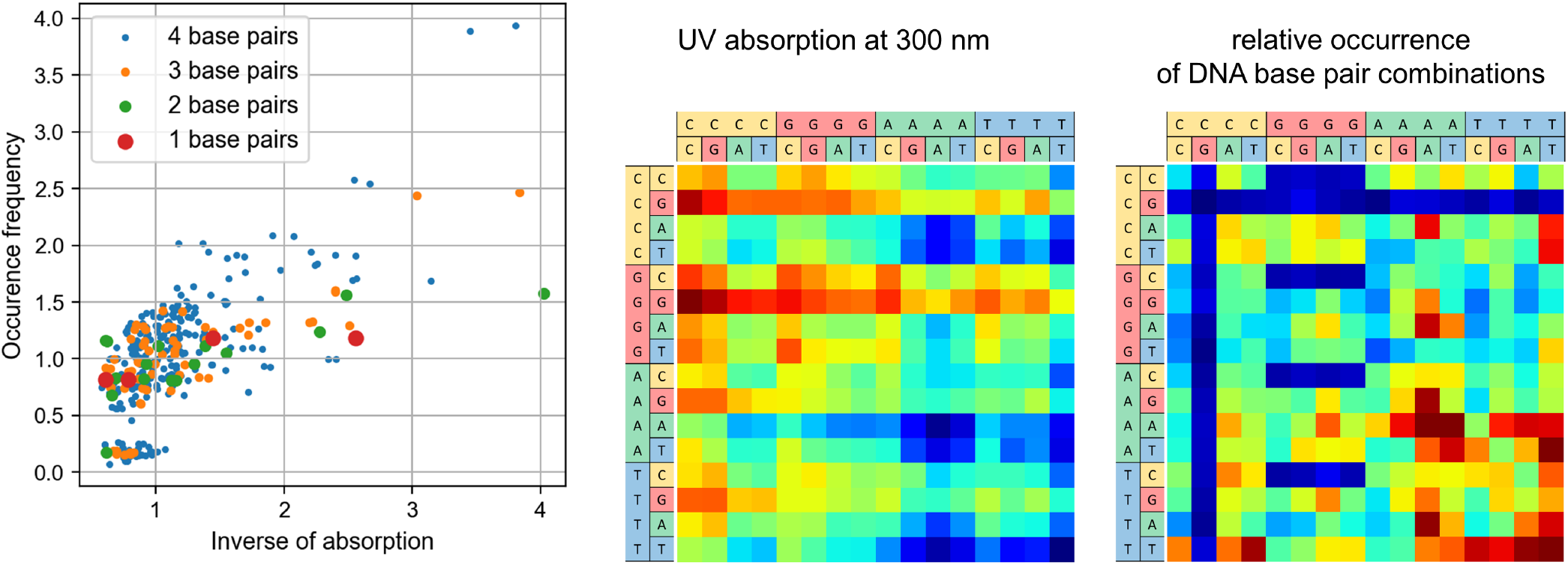
Left: The occurrence frequency of DNA monomers (red), dimers (green), trimers (yellow) and tetramers (blue) are plotted against the inverse of their absorption at 300 nm. Both quantities are normalized to have a mean of one for each group. Middle/right: UV absorption at 300nm and relative occurrence of all 256 DNA tetramers, each visualized as a single square. Blue corresponds to low absorption/low occurrence and red to high absorption/high occurrence. Occurrence is truncated at 5 *·* 10^7^ for visualization purposes due to exceptional high occurrence of some A/T-rich sequences.

The scatter plot in Fig. 2 exhibits a distinct cluster near the origin. The sequences belonging to this cluster all contain the dimer CG, which is also prevalent in the nullomer sequences. CG occurs only rarely in genomes because CpG sites are associated with elevated mutation rates: methylated cytosine can undergo deamination to thymine, resulting in a C *→* T transition [25]. The depletion of CG motifs is likewise evident from the red bars in Fig. 2 (right).

Finally, we investigate the spatial origin of UV absorption within the DNA sample. This analysis reveals whether the absorption is dominated by the nucleobases or the backbone and, for longer sequences, identifies the nucleotides that contribute most strongly to the absorption process.

To identify the microscopic origin of the absorption, we Fourier transform the time-dependent dipole moments associated with the individual basis functions and extract their contributions to the total absorption spectrum. By assigning these contributions to the corresponding atoms and removing the constant background, we obtain a spatially resolved picture of the atomic contributions to the absorption at a given wavelength.

Figure 3 illustrates the spatially resolved absorption at 300 nm for the DNA dimer TG. The analysis shows that guanine and cytosine contribute more strongly to the absorption than thymine and adenine. The same method is applied to the first nullomer sequence, whose absorption pattern is also visualized in Fig. 3. From the top-down view, it becomes apparent that the absorption is dominated by the nucleobases rather than the DNA backbone. While almost all CG base pairs contribute significantly to the absorption, three adenine nucleotides also exhibit substantial contributions. These observations indicate that the absorption of longer DNA sequences cannot be explained solely by the properties of the individual nucleotides, but instead reflects a more complex collective response of the molecular structure.

**FIG. 3.**
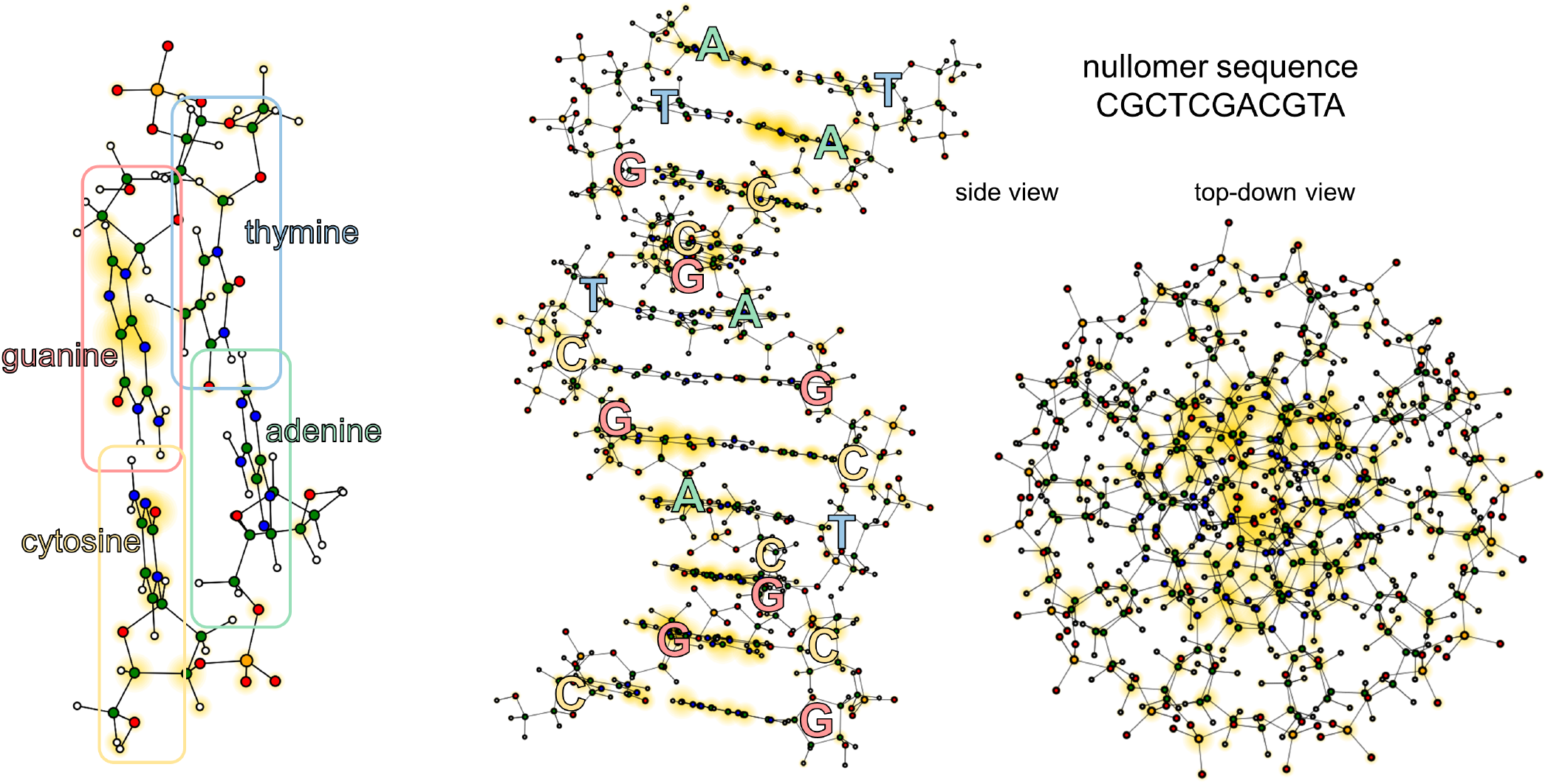
Relevant absorption regions of a DNA dimer (TG, left) and the first nullomer sample (CGCTCGACGTA, middle and right, in different perspectives). Atoms participating in the absorption at 300 nm are highlighted with yellow circles with the size of the circles proportional to their contribution.

Across all samples considered, a clear trend emerges: cytosine- and guanine-rich DNA exhibits enhanced absorption in the UVB region near 300 nm, whereas adenine- and thymine-rich DNA generally shows lower absorption. This explains the negative correlation with the occurrence frequencies in the human genome which, similar to many genomes, favors A- and T-containing sequences. Several mechanisms have been proposed to explain the relatively low GC content of the human genome [25–27], including mutation biases toward adenine and thymine [28] and the deamination of cytosine [29].

The increased absorption of nullomers can also be explained with similar reasoning. As they contain the motif CG two or three times per sequence they exhibit much higher GC content than the average of the human genome. Consequently, their absorption in the UVB region is higher than for random or synonym sequences.

Interestingly, the most common UV-induced damage is the formation of the cyclobutane pyrimidine dimer T=T [4, 30], despite it having very low absorption itself. This is partially explained by its high quantum damage yield [4] which can overcompensate the low absorption (see Supplementary information, Figure 8 for a comparison with Kufner et al.). Additionally, it should be considered that light absorption in DNA is not localized to a certain nucleobase as excited states span multiple base pairs [31]. Consequently, it is possible that absorptions happens via a larger sequence of base pairs of which some are not directly involved in the potential damages that can occur.

Spectroscopic data for single nucleobases shows, by a large margin, the highest absorption for guanine, followed by cytosine, thymine, and then adenine [32]. Additionally, cytosine and guanosine phosphates exhibit larger absorption than adenine and thymine phosphates [33].

Matallana-Surget *et al*. postulated on the basis of experimental observations that GC-rich genomes exhibit an increased susceptibility to UV-induced mutations [34]. Consistent with our findings, Sutherland *et al*. reported a correlation between genomic GC content and DNA absorption in different species [35]. These observations are in good agreement with the sequence-resolved theoretical analysis presented here.

In summary, we have performed a systematic theoretical investigation of UV absorption over a broad range of DNA sequences and identified a pronounced dependence on base composition, with AT-rich sequences exhibiting substantially lower UVB absorption than GC-rich sequences. This leads to lower absorption of UVB radiation in genomes with a lower GC content, such as the human genome, compared to random DNA sequences.

More generally, the presented framework enables fully quantum-mechanical investigations of electronic processes in large nucleic-acid systems and opens the door to studies of RNA, DNA–ligand interactions, excited-state dynamics, and sequence-dependent photostability at unprecedented scales.

## Acknowledgements

—This work was supported by the Deutsche Forschungsgemeinschaft (DFG) through RTG 2749/1, “Biological Clocks on Multiple Time Scales”. Both authors would also like to acknowledge networking support by the COST Action CA21169, DYNALIFE, supported by COST (European Cooperation in Science and Technology).

The authors acknowledge computing time provided on the high-performance computer Lichtenberg II at TU Darmstadt, part of the National High Performance Computing Center for Computational Engineering Science (NHR4CES), as well as computational resources provided by the University of Kassel and the authors’ research group.

Biomolecular graphics were generated with UCSF ChimeraX [19–21].

## Data availability

—The source code used for the RT-TDHF calculations is publicly available at Zenodo (https://doi.org/10.5281/zenodo.16889519).

## Notes

### Competing Interest Statement

The authors have declared no competing interest.

### Summary of Updates

List of changes for the second version: - The focus of the manuscript has been changed from DNA damage due to UV radiation to DNA absorption of UV radiation. This includes changes to the title, the abstract, the introduction, the discussion and the summary of the manuscript. - Addition and discussion of literature regarding sequence-dependent UV damage, computational spectra (in the introduction) and further experimental absorption studies (in the discussion). - Control calculations with time-dependent density functional theory (TDDFT) at a high accuracy have been made for DNA monomers, dimers, single nucleobases and nucleotides. Additionally, for DNA dimers, calculations with water as an environment as well as with water and Magnesium ions were done. These correspond to the supplementary material section "Electronic structure benchmarks and environment effects with TDDFT" and Supplementary Figures 2-7. - Sensitivity analyses with respect to the wavelength correction and the corresponding tetramer mean spectrum calculation have been performed. These correspond to the supplementary material sections "Sensitivity analysis of the wavelength correction" and "Sensitivity analysis of the mean calculation" as well as Supplementary Figures 8 and 9. - A comparison with tetramer damage yields by Kufner et al. was added. This corresponds to the supplementary material section "Comparison with DNA damage yields by Kufner et al." and Supplementary Figure 10. - An atom involvement analysis in the absorption for selected tetramer sequences has been done. This corresponds to the supplementary material section "Intensities per atom for selected tetramers" and Supplementary Figure 11. - Additional information and sequences for random DNA samples used in this work are provided. The corresponding supplementary material section is called "Used random DNA sequences and their generation". - The color coding and caption of Figure 2 has been changed. - Various small changes, i.e. references to supplementary material, rewording because of imprecise language, correction of figure numbers and addition of further clarifications (regarding absorption and absorbance, normalization in Figure 2 and the used computational framework).

