## Supplementary material for "Linking UV absorption of DNA with base pair sequences"

### Supplementary material for
### Linking UV absorption of DNA with base pair sequences

Luc Wieners and Martin E. Garcia
*Institute of Physics, University of Kassel, 34132 Kassel, Germany*

*Structure visualization*—The different types of studied DNA samples are visualized at an atomistic level in Figure 1 below.

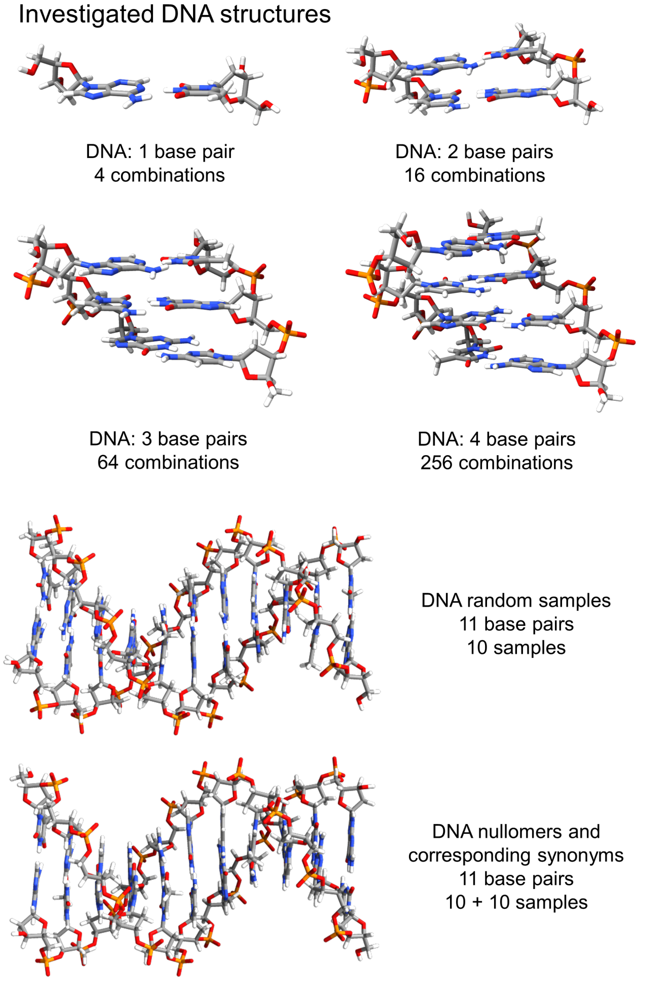


FIG. 1. Atomistic visualization of the studied DNA samples. Top row: Examples for DNA monomers, dimers, trimers and tetramers. Bottom row: 11-base pair DNA samples used for computing the UV absorption spectra of nullomers, synonyms and random DNA. One nullomer and one random sample are depicted, only subtle differences can be seen due to the different base pair sequences. Visualization with UCSF ChimeraX [1–3].

*Technical information about RT-TDHF*—A real-time approach is used by us as it scales more favorable than linear-response formalisms for large-scale systems. RT-TDHF has a bottleneck regarding its scaling in the form of various matrix operations at an $\mathcal{O}(n^3)$-cost. The theoretical $\mathcal{O}(n^4)$-scaling of the Hartree-Fock method, caused by the electron repulsion integrals, can be reduced to $\mathcal{O}(n^2)$ or $\mathcal{O}(n)$.

In the RT-TDHF calculations, the system is excited with a small initial electric field pulse and then propagated over 500 atomic units of time (12.1 fs). The propagation is done with a predictor-corrector scheme [4] and runs over 2000 steps. The dipole moments $d$ are calculated at each step and saved. To compute the absorption, a Fourier transform of the dipole moments is taken. Then, the absorption corresponds to the imaginary part of this Fourier transform divided by a Fourier transform of the electric field pulse, multiplied with the frequency $\omega$ and summed over all three spatial dimensions:

$$A(\omega) = \sum_{\alpha=x,y,z} -\frac{4\pi\omega}{3c} \Im \left( \frac{\int_0^{t_{\max}} e^{i\omega t} d_\alpha(t) e^{-\gamma t^2} \, dt}{\int_0^{t_{\max}} e^{i\omega t} E_\alpha(t) \, dt} \right), \tag{1}$$

with an attenuation factor $\gamma = 1.5 \cdot 10^{-4}$ a.u. and the speed of light $c$, the integral is then numerically evaluated with the data obtained for the time steps. Note that an attenuation factor in the form of an exponential function is necessary since we only have a finite signal in the time domain. Nonphysical oscillations are therefore suppressed, often a decreasing exponential is used. Here, we use a Gaussian function since the resulting signal in the frequency domain then decreases faster which suits the behavior of the DNA spectra in the UVB region better.

*Used random DNA sequences and their generation*—For the calculation of the UV absorption of longer DNA sequences, random DNA sequences were used. These were generated in a way that their distribution of nucleotides is roughly equal (deviations within 1%) to 30% adenine, 30% thymine, 20% cytosine, and 20% guanine. For each of the ten sequences with eleven base pairs, each subsequent base pair was chosen with the probability above irrespective of previous base pairs or sequences. Since only 110 base pairs are generated, the deviations often cause the global distribution of nucleotides to deviate from the intended distribution. We therefore generated multiple sets of random sequences and chose the first one that globally fulfilled the nucleotide distribution to the demanded accuracy.

The ten used DNA sequences are: ATCCATACTCA, GTGGACGCTTT, ATTATAGCTCT, GGGAATCAA-TA, ACTAACTAGGA, TCTTATGTTTG, AGAGACA-TTCT, CGCGCGTACAA, CTCGGCCATAA, AGATG-

ATATTG.

*Electronic structure benchmarks and environment effects with TDDFT*—Since the used RT-TDHF framework has limitations regarding its accuracy due to basis set size, long range interaction treatment and neglect of correlation, control calculations were performed for DNA dimers and monomers. To do so, we used the quantum chemistry program ORCA [5–21]. The control calculations used linear-response time-dependent density functional theory (LR-TDDFT) [22] with the Tamm–Dancoff approximation (TDA) [23], the def2-TZVP basis set [24], the CAM-B3LYP [25] exchange-correlation functional, resolution-of-the-identity [6, 7, 16] Coulomb integrals and chain-of-spheres exchange (RIJCOSX) [8, 10, 15, 16, 18] as well as the Grimme D4 dispersion correction [26] for preparation calculations.

In addition to computing high-accuracy TDDFT spectra for validation purposes, the influences of water as a solvent and Magnesium ions were investigated. Therefore, the absorption spectra of the DNA dimers were computed without environment, with water as a solvent and with water and one $Mg^{2+}$ ion per dimer. For modelling the solvent, the conductor-like polarizable continuum model (CPCM) implemented in ORCA was used, and Magnesium ions were added using UCSF Chimera [2]. The geometry of the systems was optimized using the same settings in combination with the D4 dispersion correction. The dimers with and without water were optimized over 40 steps and the dimers with ions over 60 steps, all other systems were optimized until convergence.

The discrete excitation energies of 30 computed roots from LR-TDDFT were visualized using both Gaussian (0.3 eV standard deviation) as well as Lorentzian profiles (0.3 eV half-width at half-maximum). Consequently, the representation is different to real-time methods such as RT-TDHF which do not give a stick spectrum as an output. The wavelength correction method used for RT-TDHF is not utilized as it is a different method and since in the high-accuracy calculations only the low-energy part of the spectrum is resolved, a re-fitting of shift and scaling factor would not be sensible. We therefore use the uncorrected spectrum at 300 nm for comparisons and additionally provide correlation values dependent on the wavelength used for comparison.

Investigations have been done for all dimers (Figure 2), all dimers in water (Figure 3), all dimers in water with a magnesium ion (Figure 4), all monomers in water (Figure 5), and for single nucleobases (adenine, cytosine, guanine, thymine) as well as single nucleotides (dAMP, dCMP, dGMP, dTMP) in water (Figure 6). For all of these calculations, the spectra are shown between 200 nm and 320 nm as well as between 260 and 310 nm as an enlargement. Both Gaussian and Lorentzian profiles are shown. In the cases of the monomers and the dimers, the computed absorption with LR-TDDFT at 300 nm is compared with the RT-TDHF absorption at 300 nm (wavelength-corrected) as well as with the sequence counts of dimers in the human genome, both for Gaussian and Lorentzian profiles. To consider the wavelength-dependence, the correlation coefficients of all aforementioned comparisons is plotted for the TDDFT absorption values taken at a different wavelength between 260 and 310 nm.

Among all comparisons, the LR-TDDFT data correlates positively with the RT-TDHF calculations and negatively with the sequence counts. The observed differences can be explained by the different methods and their different accuracies as well as different environments. It can be observed that the absolute magnitude and the magnitude of the differences varies strongly between Gaussian and Lorentzian profiles due to their different nature in their decay. This indicates that a quantification of the amount of absorption is difficult, the general trend of higher absorption in GC-rich however can be observed in both methods and all environments.

If we align peak positions, the spectrums could be shifted by approximately +30 nm. It is therefore interesting to also consider the wavelength-dependent correlation values at 270 nm. These sometimes show weaker correlations but still maintain a positive correlation with the RT-TDHF values and a negative correlation with the dimer occurrences for all four sets of investigated molecules and for both Gaussian and Lorentzian profiles.

Conformational sampling was investigated with the four single nucleotides dAMP, dCMP, dGMP and dTMP. Ten conformers provided by each corresponding PubChem [27] entry of the four discussed molecules were investigated. The results are visualized in Figure 7. While conformers show slightly different absorption, the different molecules can still be distinguished in most cases and the described trends are still apparent if the mean of ten conformers instead of an idealized structure is considered. It should be noted that DNA has less conformational freedom than the single nucleotides studied here.

*Sensitivity analysis of the wavelength correction*—The wavelength correction equation $\lambda_{\text{corr}} = 1.1 \cdot \lambda_{\text{calc}} + 20$ nm was investigated regarding the influence of its scaling factor (1.1) and its shift (20 nm). To obtain these values, an error metric was used which considers six values for which ideal values are given. The position of the absorption maximum should be at 260 nm, the position of the absorption minimum at 230 nm. Three ratios of the absorption at fixed wavelengths should have values of $A_{260}/A_{280} \approx 1.8$, $A_{260}/A_{230} \approx 2.0$, $A_{260}/A_{300} \approx 20.0$. The differences of the actual and desired values are computed and weighted with prefactors of 0.1, 0.1, 1.0, 1.0 and 0.25, respectively. Then, these values are squared and multiplied with the mean of the squared deviation between the reference absorption curve and the actual absorption curve. This was evaluated numerically between 220 and 320 nm using a step width of 0.1 nm. The result was divided by 1,000 for normalization purposes.

Using this procedure, shifts between −20 nm and +60 nm as well as scaling factors between 0.9 and 1.3 were investigated. The resulting values are shown in Figure 8 (plotted logarithmically). Additionally, the correlation with tetramer occurrence values is computed for these parametrizations to investigate its dependence on the wavelength correction equation.

In addition to this, the observed inverse correlation between sequence absorption and sequence occurrence has been investigated for different wavelengths. The correlation between the absorption values at 300 nm and other wavelengths has been calculated as well. The results are shown in Figure 8, bottom panels.

*Sensitivity analysis of the mean calculation*—The mean of the absorption spectra of the tetramers is used for the calculation of the wavelength correction. It can be computed without normalization or it can be weighted with the occurrence numbers of tetramers in the human genome. Both resulting spectra are shown in Figure 9. The observed deviation is only small and therefore does not have a significant influence on the main conclusion as the wavelength correction, which is influenced by this normalization, does not alter the results significantly under small perturbations.

*Comparison with DNA damage yields by Kufner et al.*—Kufner et al. [28] measured the damage quantum yields for tetramers at 266 nm which are compared with the computed absorption of DNA tetramers using RT-TDHF at 266 nm and 300 nm in Figure 10. For the latter comparison, it has to be kept in mind that damage events can be strongly wavelength-dependent [29, 30]. Additionally, it has to be considered that Kufner et al. reports the use of single-stranded DNA wheras the computational spectra were done with double-stranded DNA. Correlations values of −0.405 and −0.443 are obtained, respectively.

This can be explained by the fact that absorption in T-containing sequences is in general lower whereas those sequences are more prone to UV-induced damages such as TT dimers. Consequently, sequence-dependent absorption does not correlate with sequence-dependent damage yields.

*Intensities per atom for selected tetramers*—The Fourier transform used for the spectra calculation based on the dipole moment calculation from RT-TDHF can be broken up into individual atoms (constant offsets are subtracted) to estimate which parts are involved in the absorption. This is done for the four sequences ATTA, CTTC, GTTG and TTTT (see Figure 11, read sequences from bottom to top), based on Kufner et al. [28] where the effects flanking bases [31] of the sequence TT were investigated. Here, we see high absorption for the sequences GTTG and CTTC in the cytosine and guanine nucleotides. Notably, GTTG has the lowest and TTTT the highest measured quantum yield, showing that sequences that absorb a lot do not suffer from a high damage rate, and vice versa. This agrees with the negative correlation observed in the previous section.

Additionally, the sequences CGCC and CCGC as well as TATA and ATAT were investigated which show differing absorption at 300 nm despite having similar structures. It can be observed that CGCC (compared to CCGC) shows more absorption involvement in most parts of its structure and in TATA the adenine nucleotides absorb more than in ATAT.

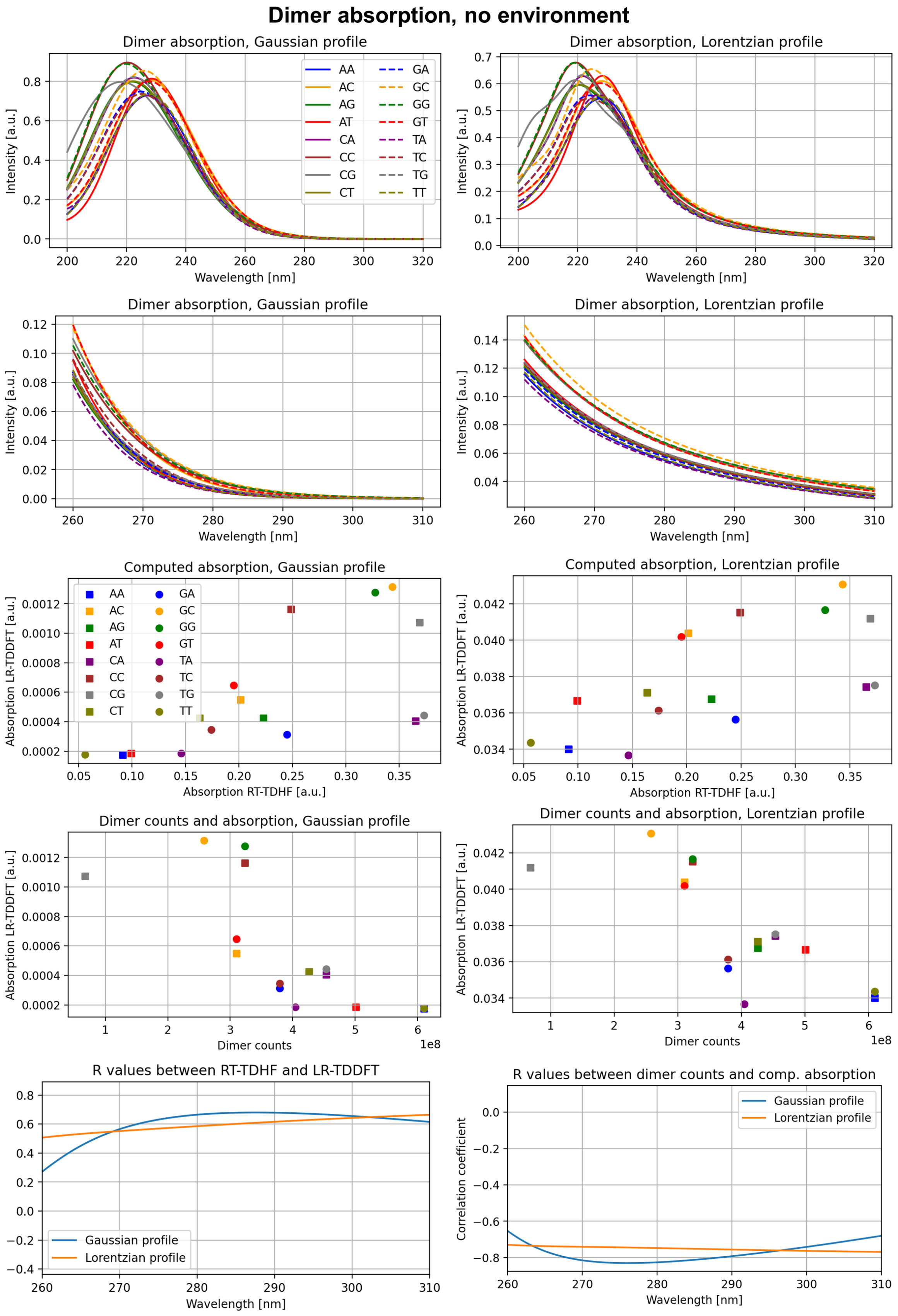


FIG. 2. Computed absorption spectra with ORCA for DNA dimers.

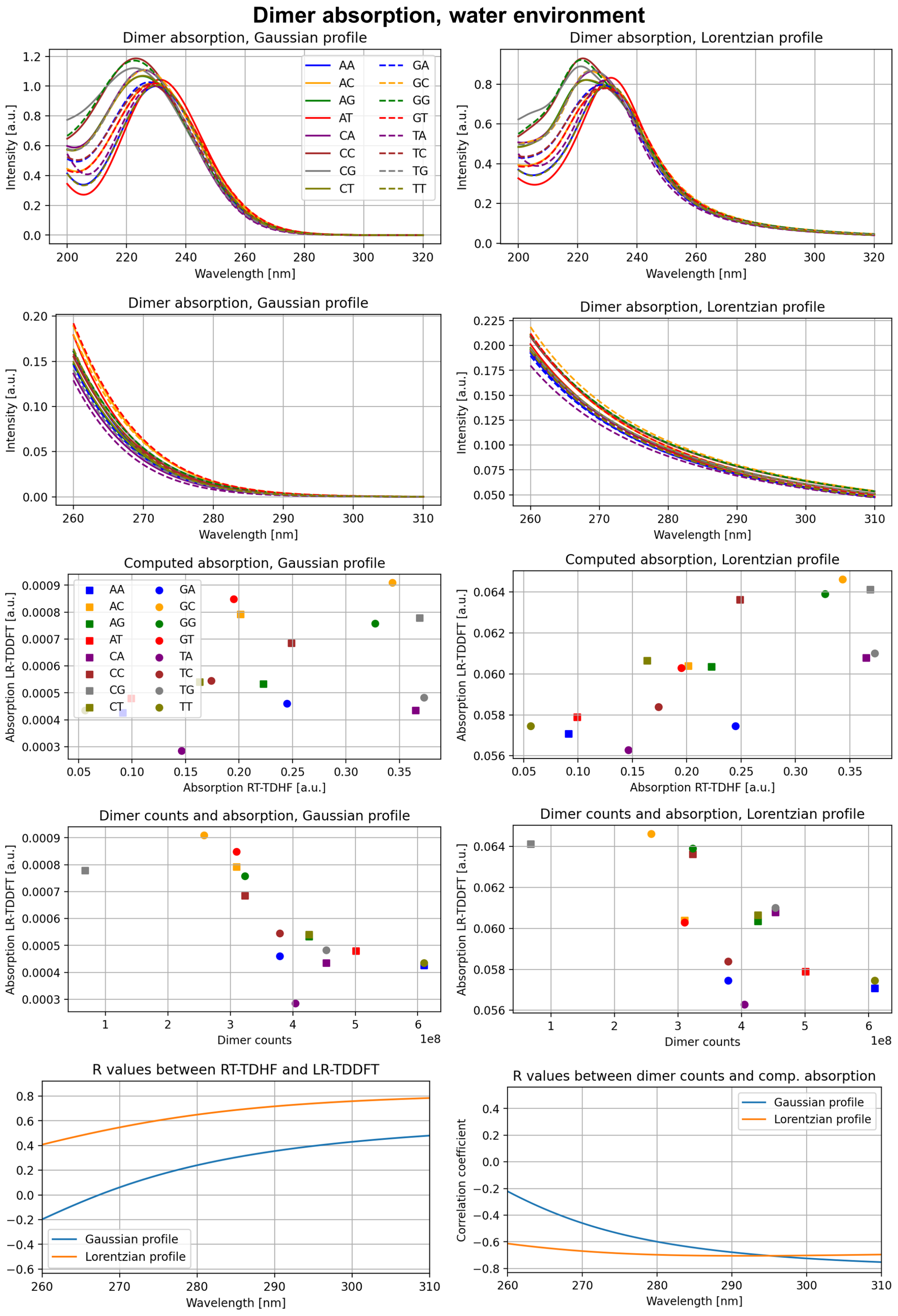


FIG. 3. Computed absorption spectra with ORCA for DNA dimers with a water environment.

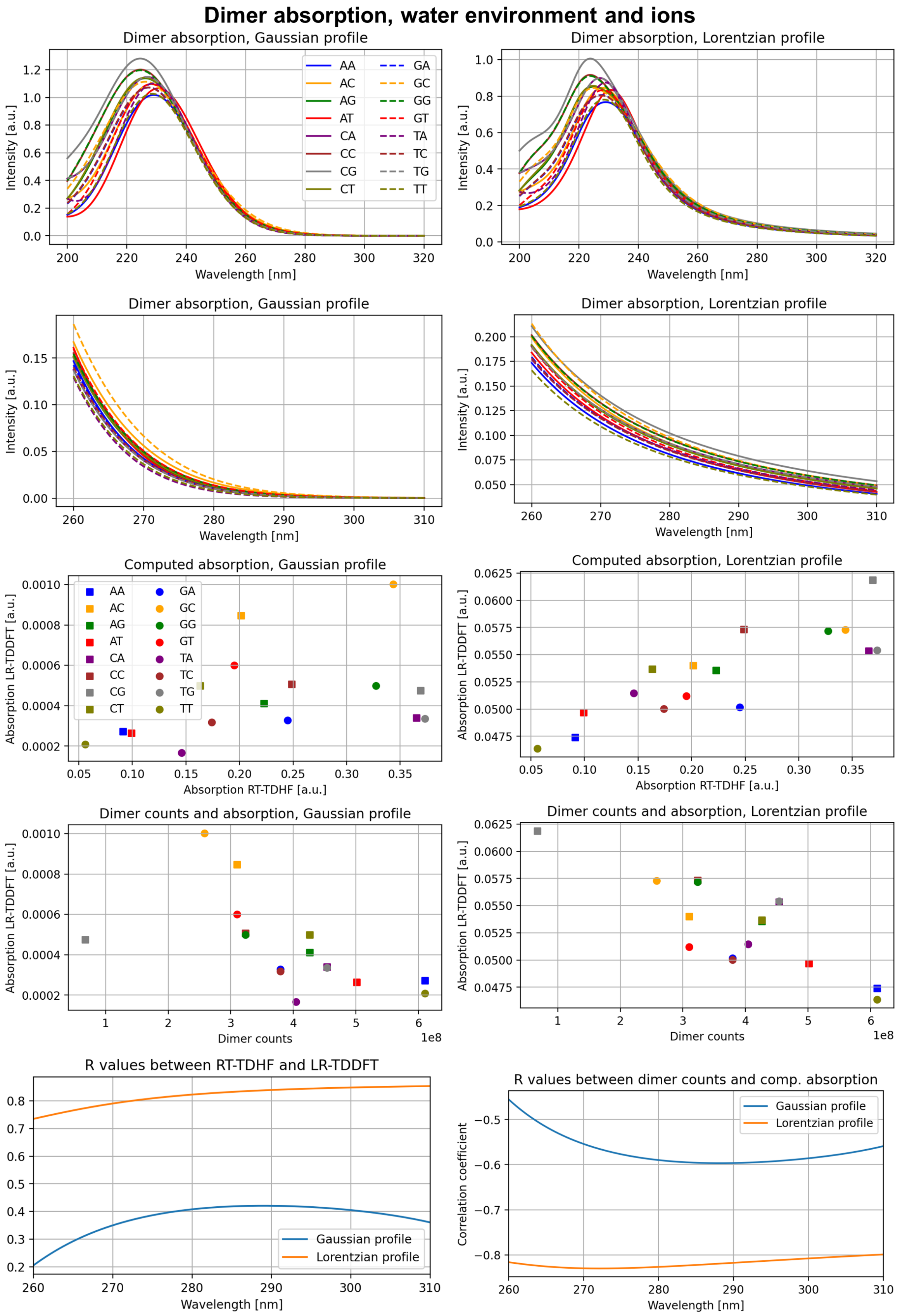


FIG. 4. Computed absorption spectra with ORCA for DNA dimers with a water environment and a Magnesium ion.

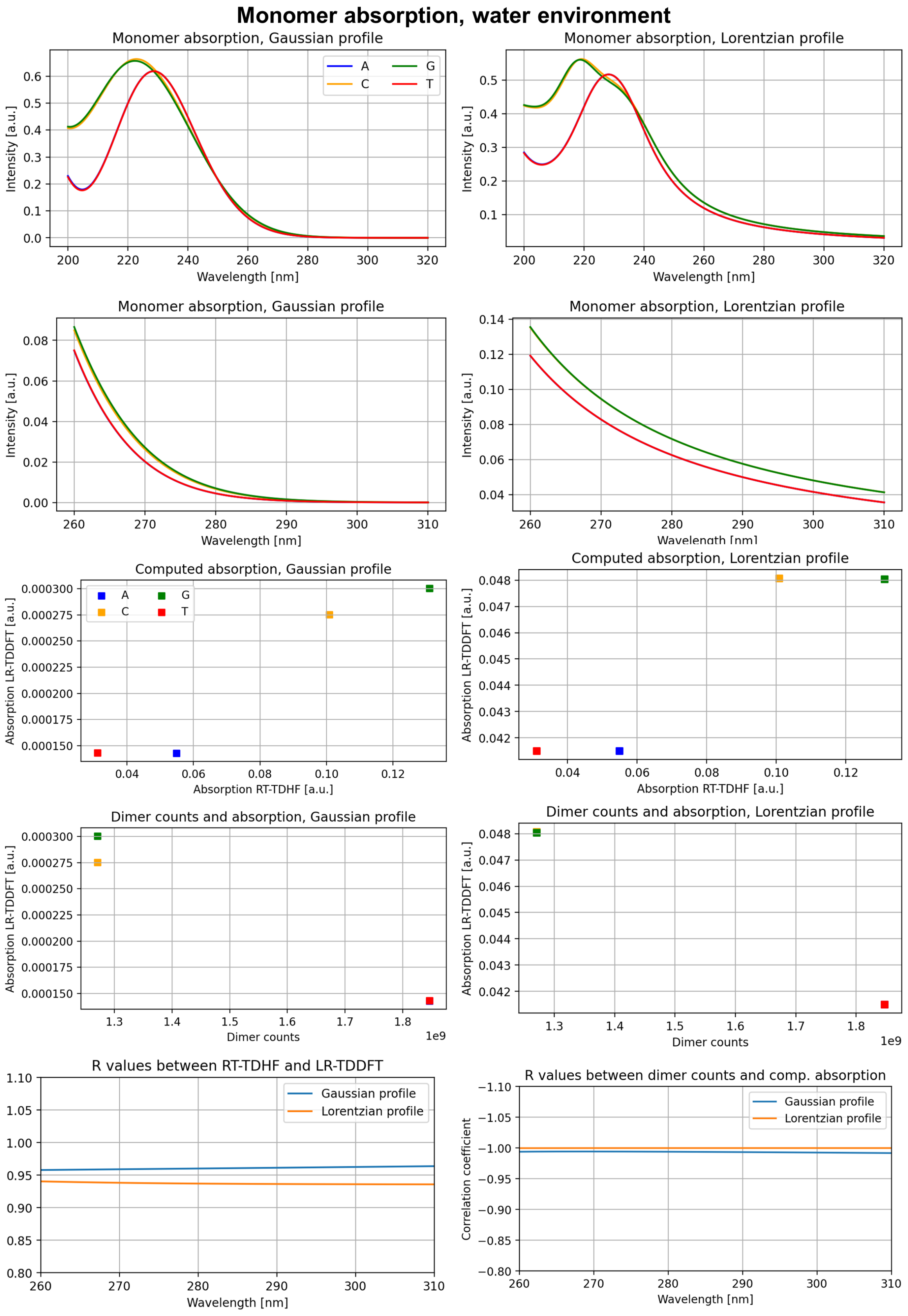


FIG. 5. Computed absorption spectra with ORCA for DNA monomers.

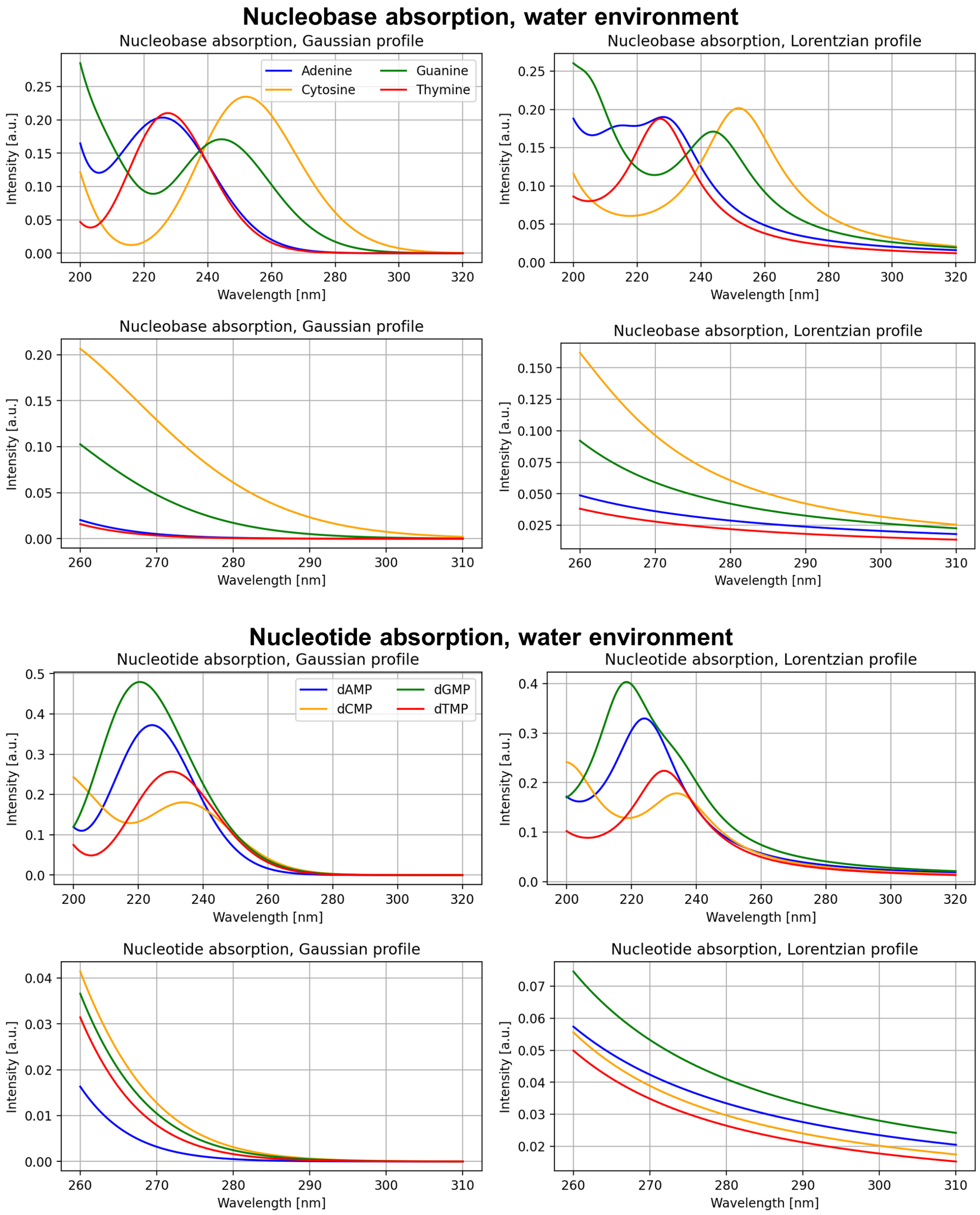


FIG. 6. Computed absorption spectra with ORCA for DNA nucleobases and nucleotides.

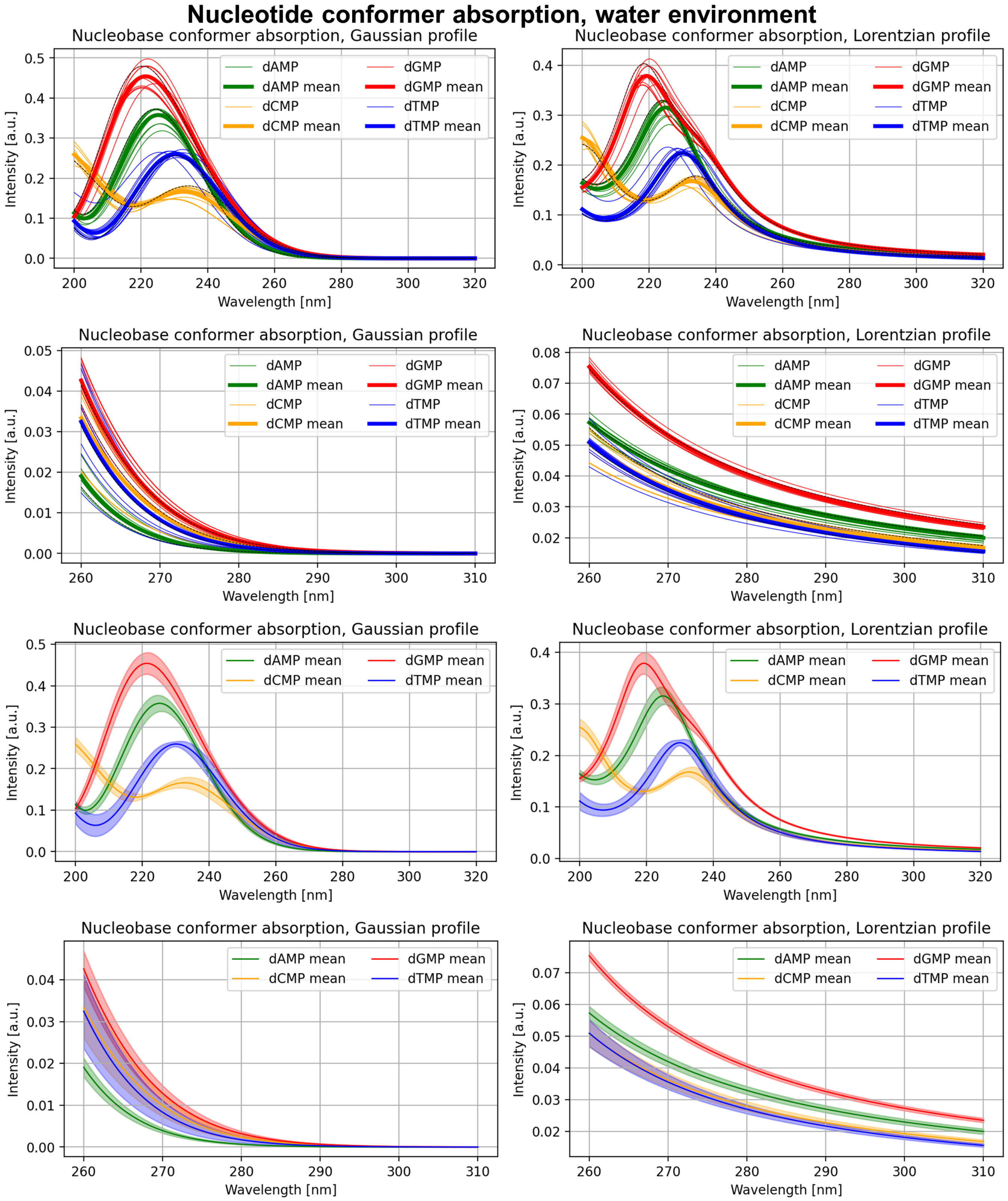


FIG. 7. Computed absorption spectra with ORCA for conformers of DNA nucleotides. In the top four panels, the conformer with the lowest energy is indicated with a dashed black line. The bottom four panels show the standard deviation of the absorption of the conformer ensembles as a colored background around the absorption mean.

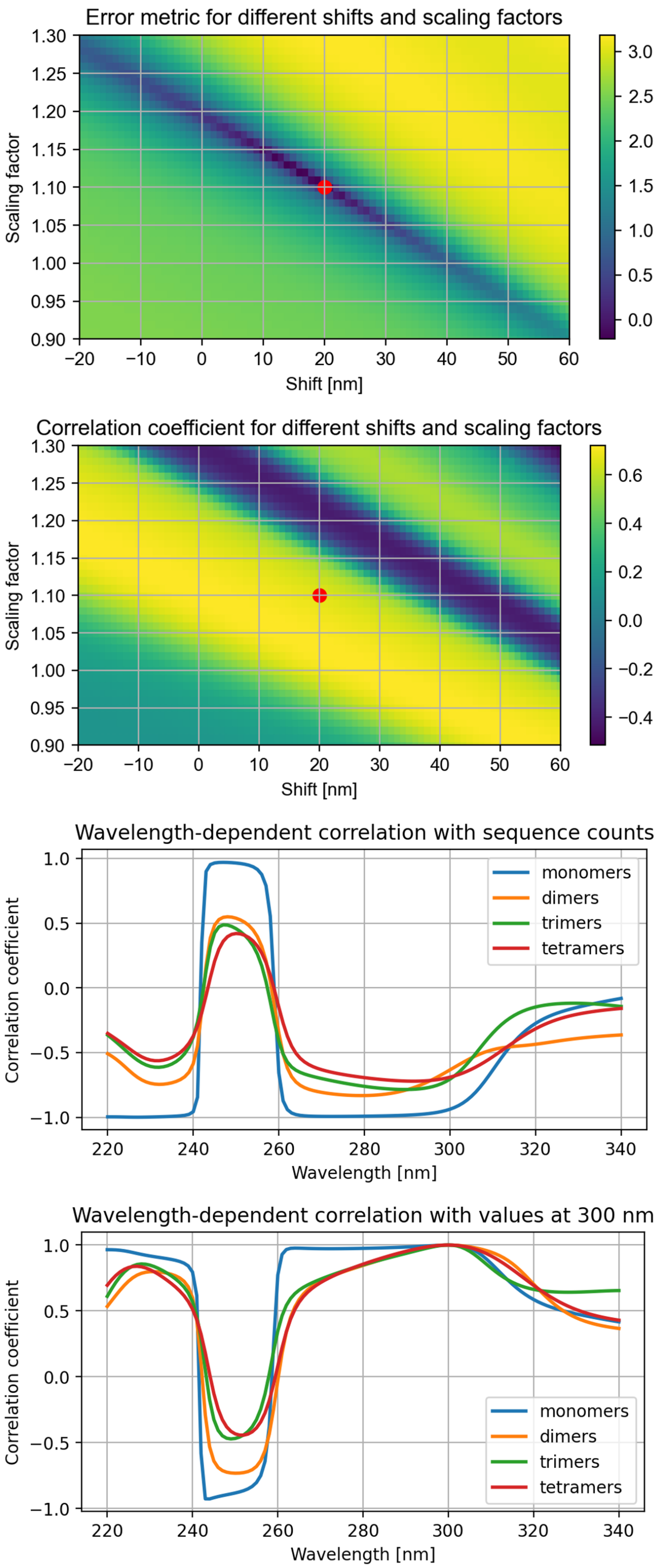


FIG. 8. Sensitivity analysis of the wavelength transformation used for spectra alignment. Top two panels: error metric and correlation coefficient (multiplied with −1) for different shifts and scaling factors, the red dot represents the used configuration. Bottom two panels: wavelength-dependent correlation with sequence counts and with values at 300 nm for monomers, dimers, trimers and tetramers.

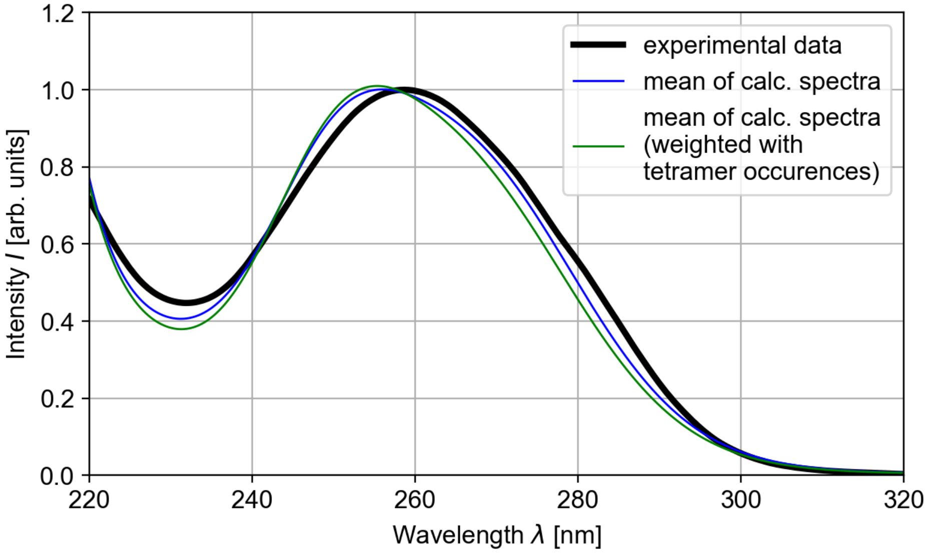


FIG. 9. The results of two different methods for the calculations of the mean of the tetramer spectra are shown: mean computation with equal weights (blue) and mean computation with weights in form of the sequence counts of tetramers in the human genome.

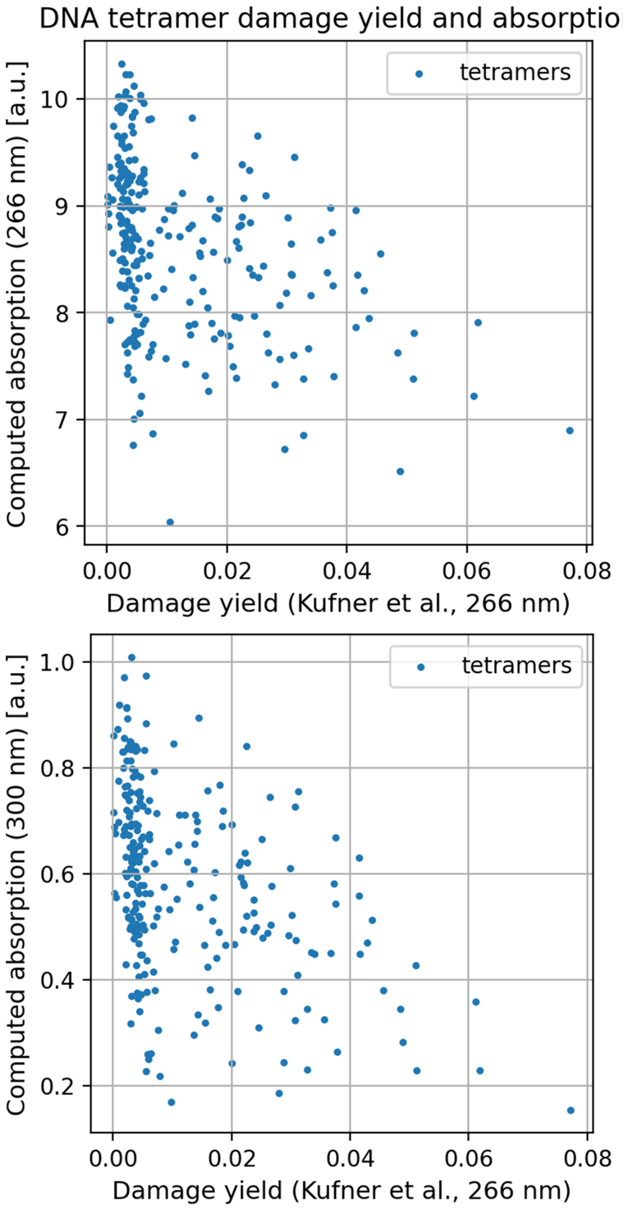


FIG. 10. Comparison of computed tetramer absorption at 266 nm (top) and 300 nm (bottom) with measured quantum damage yields at 266 nm by Kufner et al. The values are taken from Table SI-t 8, the value $\mu_{\rm olig}$ is used.

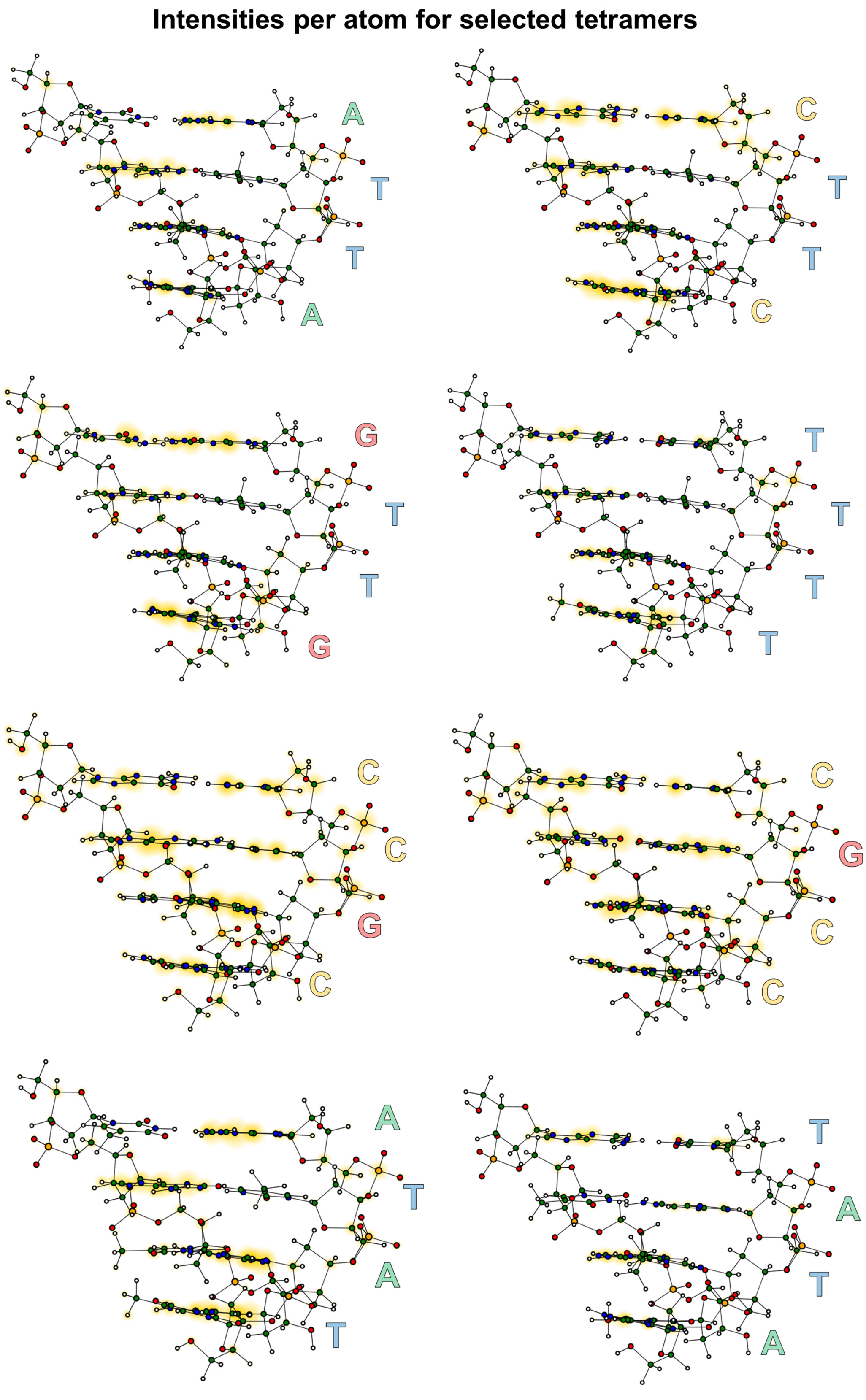


FIG. 11. Atom-dependent intensities (obtained via Fourier transforms of the dipole moments of the individual atoms) of the selected tetramers ATTA, CTTC, GTTG, TTTT, CGCC, CCGC, TATA and ATAT. Sequences are read from bottom to top.
